## Supplementary material for "Identification of a guanine-specific pocket in the protein N of SARS-CoV-2": Table 1

Table I: Data Collection and Crystallographic statistics.

**N^CTD^ in complex with GTP**

**SARS-CoV-2	N^CTD^**

### Processed data

**Crystal form I	Crystal form II**

| Beamline | XALOC (ALBA) | XALOC (ALBA) | XALOC (ALBA) |
| --- | --- | --- | --- |
| Wavelength (Å) | 0.97926 | 0.97926 | 0.97926 |
| Space group | P 1 | P 2_1_ | P 1 |
| Cell dimensions  *a, b, c* (Å) | 43.84 , 44.82 , 59.12 | 43.77 , 92.72 , 68.63 | 43.69 , 48.41 , 68.67 |
| *α, β, γ* (°) | 92.04 , 96.20 , 90.00 | 90.00 | 74.37 , 89.89 ,83.17 |
| Resolution (Å) | 44.79–1.94 | 92.72–1.80 | 66.09–2.00 |
|  | (2.04-1.94) | (1.84-1.80) | (2.05-2.00) |
| R_pim_ (%) | 0.109 (0.527) | 0.027 (0.169) | 0.052 (0.212) |
| Mean I/δ(I) | 6.2 (2) | 17.8 (4.9) | 8.9 (3) |
| CC (1/2) | 0.984 (0.704) | 0.999 (0.961) | 0.993 (0.923) |
| Unique reflections | 32080 (4753) | 50472 (2926) | 34712 (2497) |
| Completeness (%) | 96.1 (96.6) | 99.4 (97.8) | 95.5 (93.2) |
| Redundancy | 3.4 (3.5) | 6.8 (6.4) | 3.4 (3.1) |

### Refined data

| Rfactor (%) | 0.185 | 0.165 | 0.192 |
| --- | --- | --- | --- |
| R_free_ (%) | 0.239 | 0.211 | 0.245 |
| RMSD  Bond deviation (Å)  Angle deviation (º) | 0.017  1. 828 | 0.020  1. 937 | 0.014  1. 748 |
| Ramachandran Map |  |  |  |
| Favoured (%) | 96.55 | 98.14 | 97.65 |
| Allowed (%) | 3.45 | 1.86 | 2.35 |
| Disallowed region (%) | 0 | 0 | 0 |
| PDB accession code | 7O05 | 7O35 | 7O36 |

Values in parentheses correspond to the data for the highest resolution shell

R_pim_ = Σ_hkl_√(1/(n-1)) Σ_i_ | I(hkl)_i_ - <I(hkl)> | /Σ_hkl_Σ_i_ I(hkl)_i_ R_factor_= Σ‖Fo|−|Fc‖/Σ|Fo|

R_free_ is the R_factor_ calculated with 5% of the total unique reflections chosen randomly and omitted from refinement.

1
