## Supplemental Figures for "Identification of a guanine-specific pocket in the protein N of SARS-CoV-2"

### Supplementary Figure 1

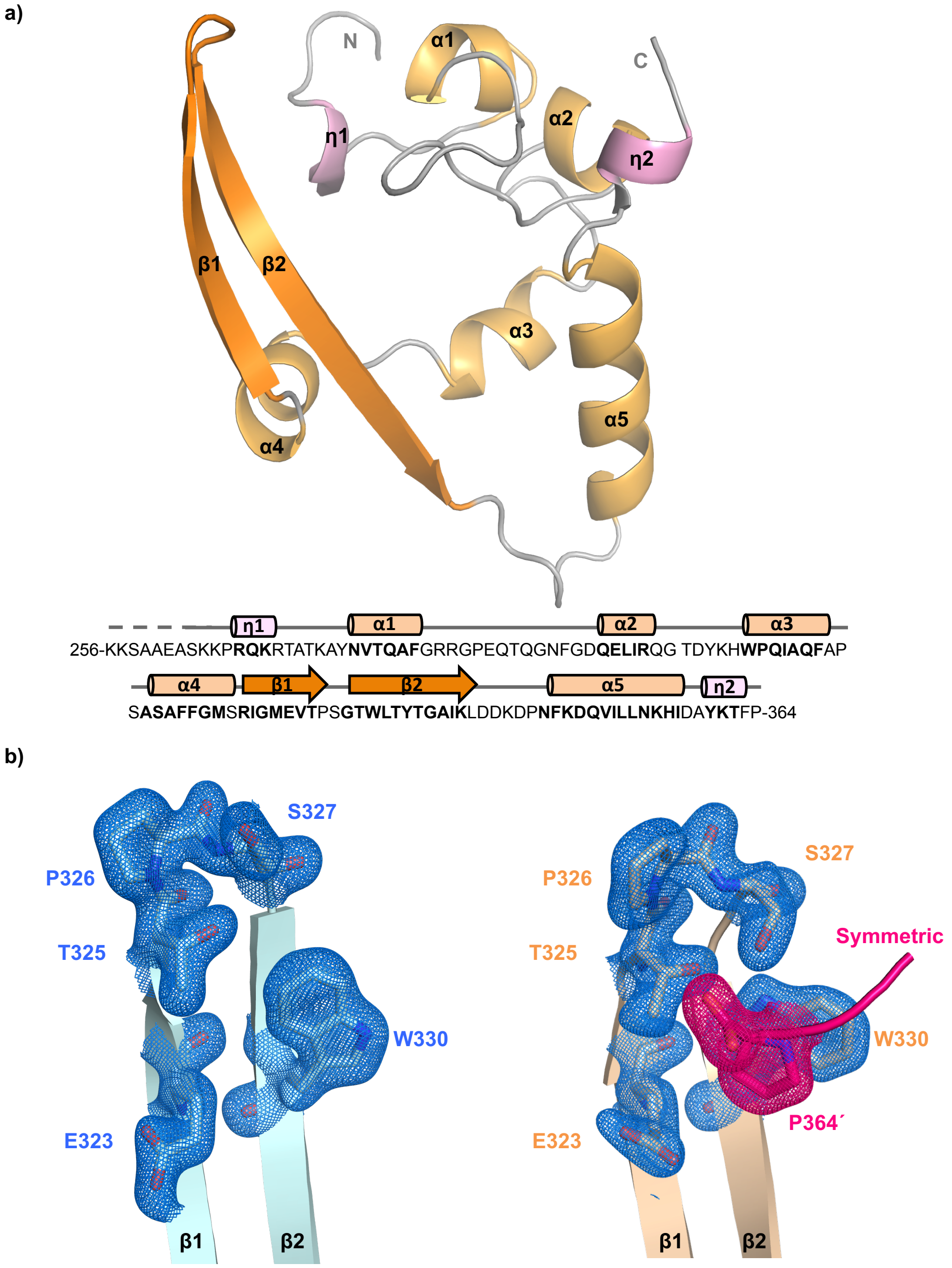


**Supplementary Fig. 1. Structure of SARS-CoV-2 N$^{CTD}$ and conformations of the β-hairpin.**
**a**) Cartoon representation of N$^{CTD}$ with secondary structural element colored in grey (loops), light orange (α-helices), dark orange (β-strands), and pink ($3_{10}$ helices). The sequence of N$^{CTD}$ is shown below the structure. **b**) Close view of the β-hairpin in the open (left; blue) and closed (right; orange) conformations. The side chains of the residues that rotate within the different confirmation are shown in sticks. The electron density map 2Fo-Fc (σ=1) is represented in blue. The symmetric molecule is coloured in magenta and the C-terminal Pro364 is shown in sticks and its electron map in magenta. Nitrogen and oxygen atoms are colored in blue and red, respectively. Secondary structural elements and residues are numbered and labeled in order from N to C terminus.

### Supplementary Figure 2

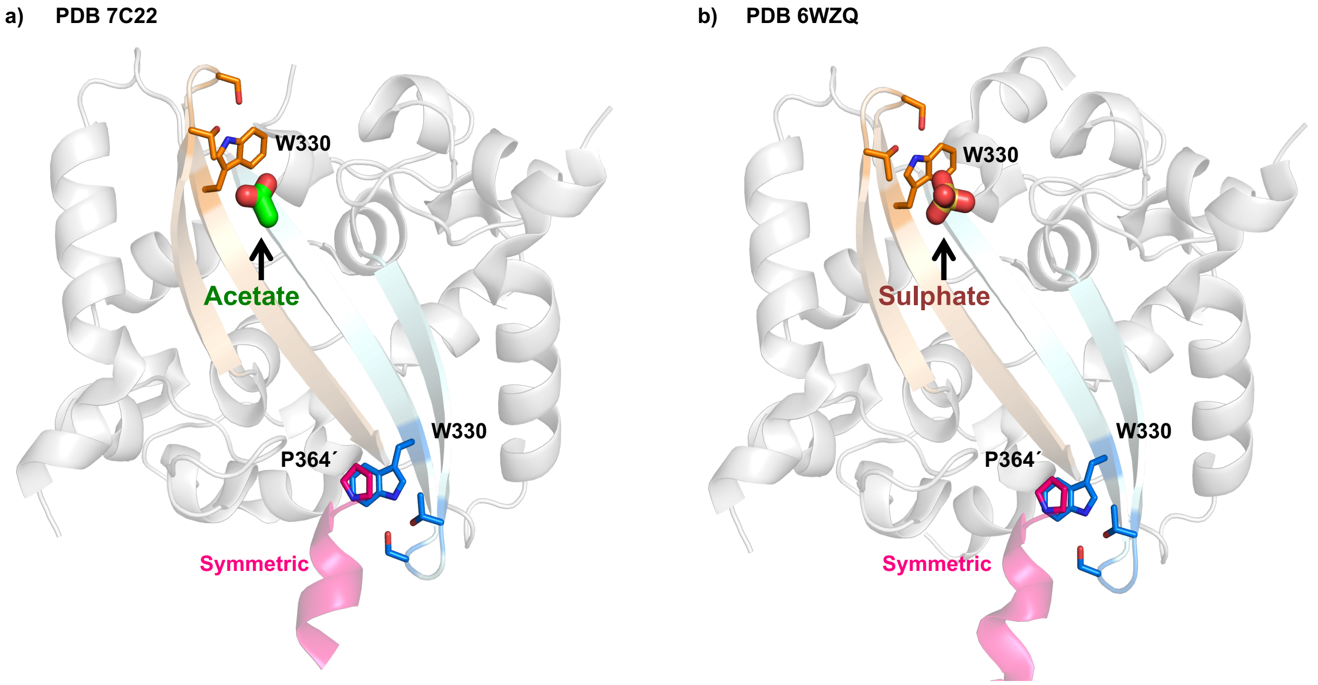


**Supplementary Figure 2**. **Acetate and Sulphate molecules bound to N$^{CTD}$ W330.**
Cartoon representation of SARS-CoV-2 N$^{CTD}$ structures PDB 7C22 (**a**) and PDB 6WZQ (**b**). The β-hairpin of each monomer is colored in pale orange and pale cyan. The helical core is coloured in white and the symmetric in magenta. Residues of the β-hairpin and the symmetric Pro364´ are shown in sticks with carbon atoms in the same colour to the subunit to which they belong. The acetate molecule is shown in sticks with carbon atom in green. The sulphate molecule is shown in sticks with sulphur atom in yellow and oxygen atoms in red.

**Supplementary Figure 3**

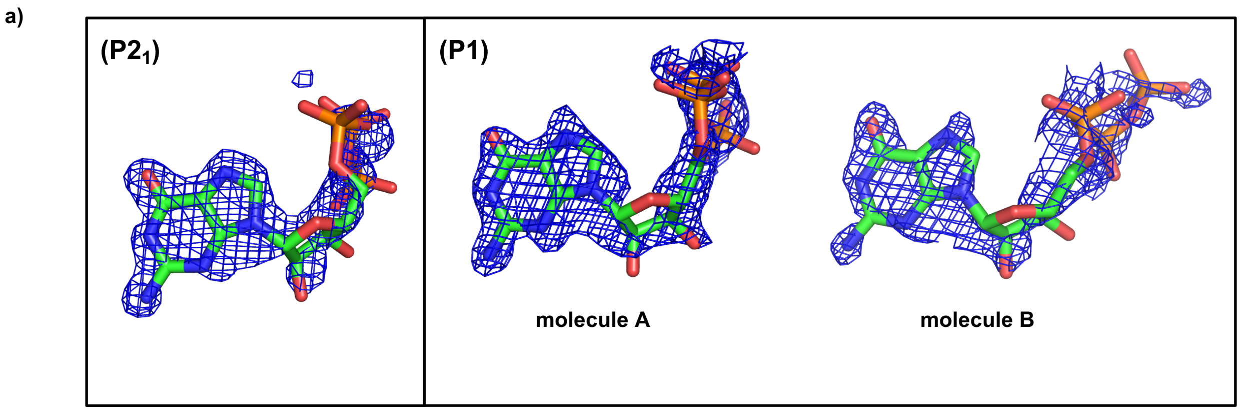


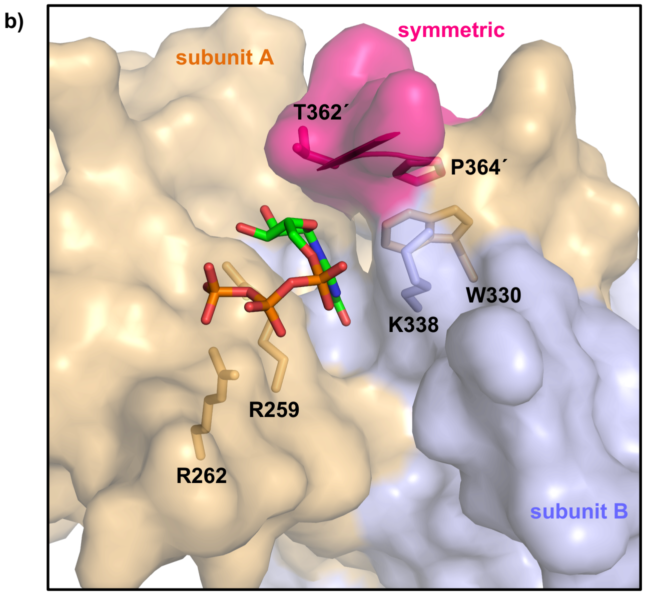


**Supplementary Figure 3. (a)** Simulated anneal omit maps of the GTP molecules. Maps are represented in blue color at σ=1, carve 1.6. **(b)** Surface representation of the GTP binding pocket. Each subunit of the dimer is colored in blue and orange, respectively, and the symmetric molecule in black. The side chains of the W330 and neighbouring residues are represented in sticks in the same colour the subunit to which they belong. The GTP is shown in sticks with carbon, nitrogen, oxygen and phosphorus atoms coloured in green, blue, red and orange colours, respectively.

**Supplementary Figure 4**

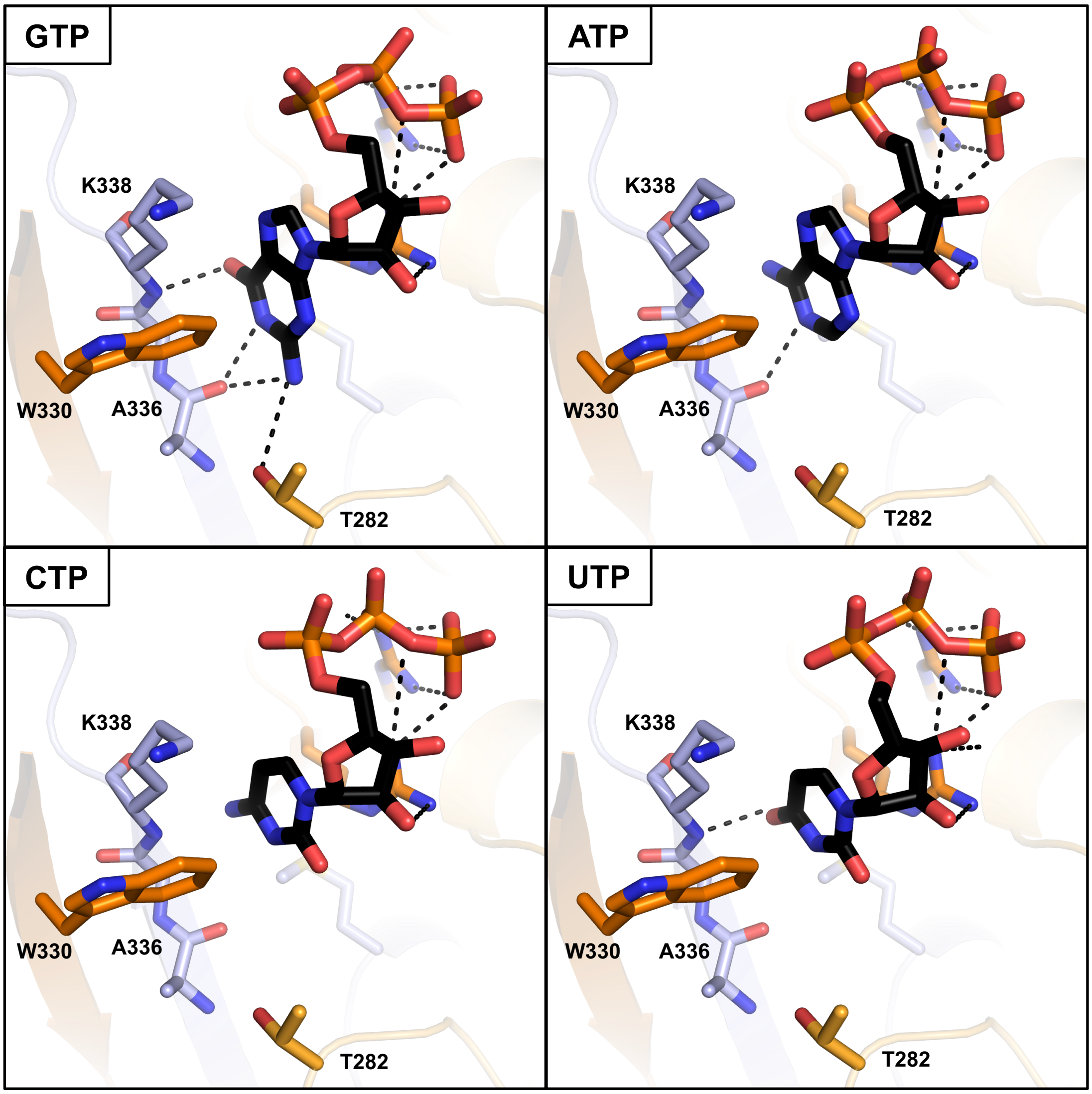


**Supplementary Figure 4. Selectivity of the GTP binding site of N$^{CTD}$ for guanine base.** The ribonucleotides ATP, CTP and UTP are superposed over the GTP molecule. The side cain of the residues that mediate interaction with GTP are shown in sticks. Structural elements and carbon atoms of each monomer of the dimer are represented in orange and blue colors.Ligands are represented in sticks, with carbon in black. Nitrogen, oxygen and phosphorous atoms are colored in blue, red and orange, respectively.

### Supplementary Figure 5

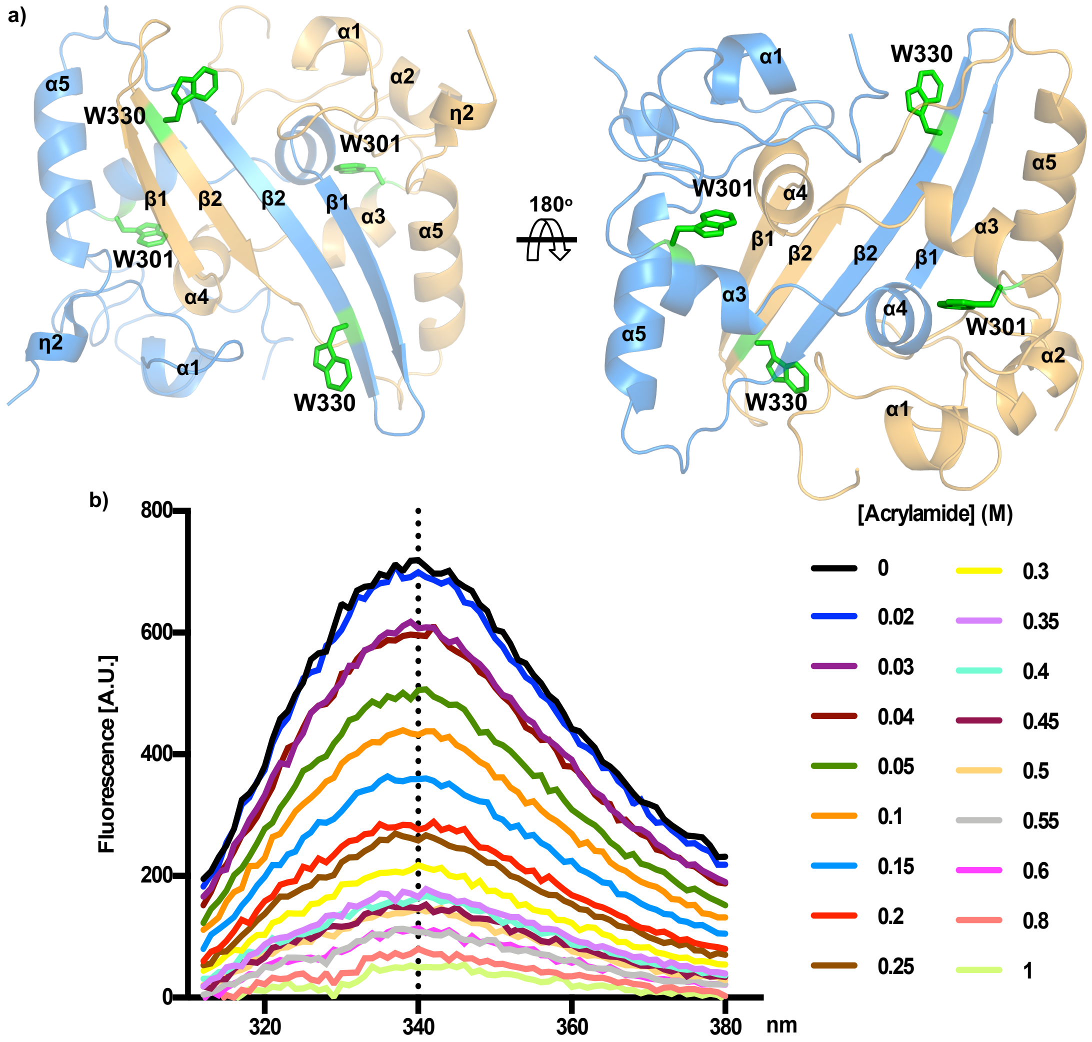


**Supplementary Figure 5**. **Tryptophan fluorescence quenching of SARS-CoV-2 $N^{CTD}$.**

**a**) Two rotated views of $N^{CTD}$ dimer. Each monomer is colored in blue and orange. Tryptophan residues are shown in sticks, labelled, and coloured in green. Secondary structural elements numbered and labeled in order from N to C terminus. **b**) Tryptophan fluorescence emission of $N^{CTD}$ from 312 to 380 nm in absence or increasing concentration of acrylamide.

### Supplementary Figure 6

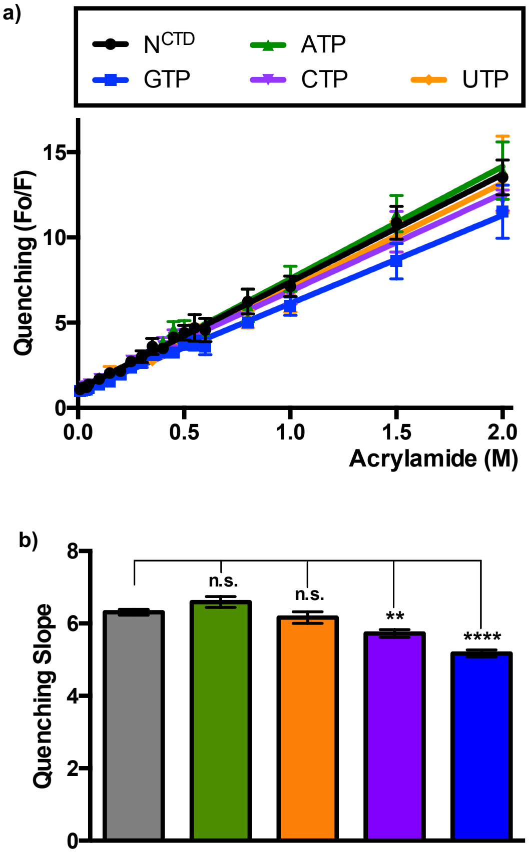


**Supplementary Figure 5**. **Interference of oxynucleotides in the quenching of the tryptophan fluorescence of SARS-CoV-2 $N^{CTD}$.**

**a)** Quenching of the tryptophan fluorescence emission at 340 nm by acrylamide in absence or presence of ATP, GTP, CTP, UTP. The quenching is represented by the ratio of fluorescence in absence of acrylamide (Fo) by the fluorescence in presence of acrylamide (F). **b)** Bar graph of the quenching slope and standard error in absence or presence of oxynucleotides. Statistical differences are indicated with asterisks above each bar (**** $p$ <0.0001, ** $p$ <0.01, n.s. $p$ >0.05).

### Supplementary Figure 7

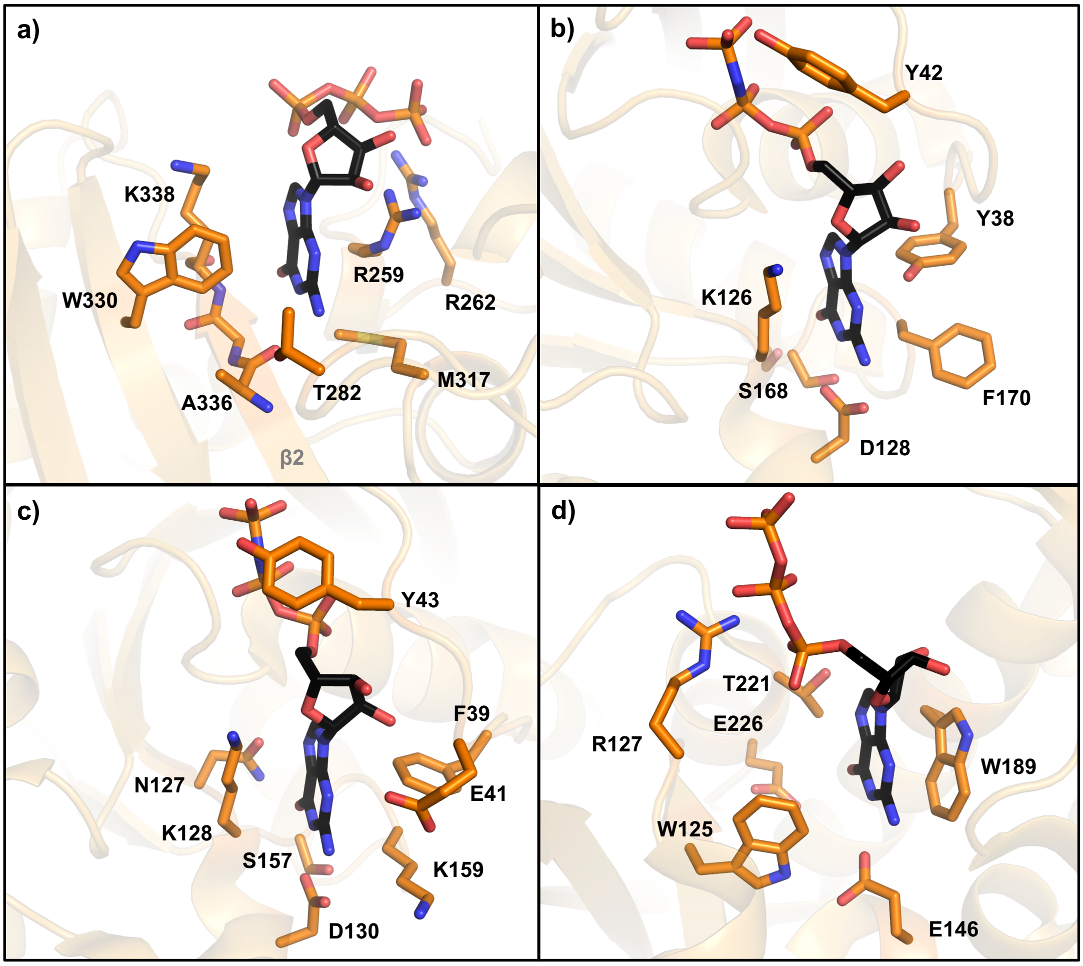


**Supplementary Figure 7. Comparatision of the GTP binding site of N$^{CTD}$ with structurally symmilar human proteins. a)** Close view of the GTP binding site of of SARS-CoV-2 N$^{CTD}$, **b)** Human Rho-related GTP-binding protein Rho6 in complex with GNP (PDB ID: 2REX), **c)** Human Ras related protein Ral-A in complex with GNP (PDB ID: 1ZC3), **d)** Human CD38 in complex with GTP (PDB ID: 3DZI). The residues that mediate interaction with the nucleotide are represented in sticks with carbon atoms coloured in orange. Ligands are represented in sticks, with carbon in black. Nitrogen, oxygen and phosphorous atoms are colored in blue, red and orange, respectively.

**Supplementary Table I: Interactions between GTP and SARS-CoV-2 $N^{CTD}$.**
Residues of $N^{CTD}$ subunits are colored in blue and orange, and the symmetric molecule in grey.

| GTP | | SARS-CoV-2 $N^{CTD}$ | | |
|---|---|---|---|---|
| **Moiety** | **Atom** | **Atom** | **Residue** | **Distance (Å)** |
| **Guanine** | **O6** | N | **K338** | 2.86 |
| | **C6** | CB | **R259** | 4.29 |
| | | CD | | 3.45 |
| | | CE | **M317** | 3.63 |
| | | CZ3 | **W330** | 3.88 |
| | | CH2 | | 4.54 |
| | | CE3 | | 4.65 |
| | | C | **A336** | 4.87 |
| | | CA | **I337** | 4.18 |
| | | C | | 4.32 |
| | | CA | **K338** | 4.35 |
| | | CB | | 3.94 |
| | **N1** | O | **A336** | 2.95 |
| | **C2** | CE | **M317** | 3.68 |
| | | CD | **R259** | 4.03 |
| | | CG2 | **T282** | 4.09 |
| | | CH2 | | 4.34 |
| | | CH2 | **W330** | 4.34 |
| | | CZ3 | | 3.67 |
| | | CE3 | | 4.64 |
| | **N2** | O | **A336** | 2.95 |
| | | O | **T282** | 3.59 |
| | **C4** | CD | **R259** | 3.63 |
| | | CZ | | 4.36 |
| | | CZ2 | **W330** | 4.99 |
| | | CZ3 | | 3.85 |
| | | CH2 | | 3.80 |
| | **C5** | CB | **M317** | 4.88 |
| | | CD | **K338** | 4.87 |
| | | CB | | 3.93 |
| | | CZ3 | **W330** | 3.96 |
| | | CH2 | | 4.15 |
| | | CG | **R259** | 4.55 |
| | | CD | | 3.34 |
| | | CZ | | 4.79 |
| | | CB | | 4.50 |
| | **C8** | CD | **K338** | 4.42 |
| | | CB | | 4.60 |
| | | CZ3 | **W330** | 5.03 |
| | | CH2 | | 4.60 |
| | | CD | **R259** | 4.23 |
| | | CZ | | 4.84 |
| **Ribose** | C1 | C | **T362** | 3.81 |
| | | CA | | 3.56 |
| | | CB | | 3.69 |
| | | CG2 | | 4.30 |
| | | CH2 | **W330** | 4.66 |
| | | CD | **R259** | 5.06 |
| | | CZ | | 4.60 |
| | C2 | CD | | 4.56 |
| | | CZ | | 3.44 |
| | O2 | NE | | 4.53 |
| | | NH1 | | 3.65 |
| | | NH2 | | 4.37 |
| | C3 | CZ | | 3.76 |
| | | CG2 | **T362** | 4.34 |
| | O3 | NH1 | **R259** | 3.82 |
| | | NH2 | | 3.52 |
| | C4 | CZ | | 5.4 |

**Conformation A**

| GTP | | SARS-CoV-2 $N^{CTD}$ | | |
|---|---|---|---|---|
| **Moiety** | **Atom** | **Atom** | **Residue** | **Distance (Å)** |
| **Pα** | O1A | NZ | **K338** | 4.22 |
| | O2A | | | 4.74 |
| **Pβ** | O1B | NH1 | **R262** | 4.89 |
| | | NH2 | | 3.39 |
| | O2B | NH2 | **R262** | 4.65 |
| | O3B | NH2 | **R259** | 3.58 |
| | | NH1 | **R262** | 4.26 |
| | | NH2 | | 4.07 |
| **Pγ** | O1G | NH2 | **R259** | 4.64 |
| | | NE | **R262** | 4.97 |
| | | NH1 | | 3.40 |
| | | NH2 | | 3.22 |
| | O2G | NH2 | **R259** | 4.85 |
| | OG3 | NH1 | | 3.98 |
| | | NH2 | | 2.67 |
| | | NH1 | **R262** | 3.32 |
| | | NH2 | | 4.42 |

**Conformation B**

| GTP | | SARS-CoV-2 $N^{CTD}$ | | |
|---|---|---|---|---|
| **Moiety** | **Atom** | **Atom** | **Residue** | **Distance (Å)** |
| **Pα** | O1A | NH2 | **R259** | 4.95 |
| | O3A | NH2 | | 3.80 |
| | | NH2 | **R262** | 4.04 |
| **Pβ** | O1B | NH2 | **R259** | 4.25 |
| | O2B | NH2 | | 3.47 |
| | | NE | **R262** | 4.54 |
| | | NH1 | | 2.62 |
| | | NH2 | | 3.02 |
| **Pγ** | O1G | NH2 | | 3.52 |
| | | NE | | 4.97 |
| | | NH1 | | 3.40 |
| | | NH2 | | 3.22 |
| | OG3 | NH1 | | 4.15 |
| | | NH2 | | 3.34 |

**Supplementary Table II: Linear regression of quenching.**

| Linear regression | $N^{CTD}$ | $N^{CTD}$+GTP | $N^{CTD-W330A}$ | $N^{CTD-W330A}$+GTP |
|---|---|---|---|---|
| **Best-fit values** | | | | |
| Slope | 6,310 ± 0,08360 | 5,173 ± 0,09601 | 1,990 ± 0,06165 | 1,738 ± 0,05859 |
| Y-intercept when X=0.0 | 1,077 ± 0,05585 | 0,9515 ± 0,06145 | 0,8802 ± 0,03946 | 1,040 ± 0,03750 |
| X-intercept when Y=0.0 | -0,1708 | -0,1839 | -0,4422 | -0,5984 |
| 1/slope | 0,1585 | 0,1933 | 0,5024 | 0,5755 |
| **95% Confidence Intervals** | | | | |
| Slope | 6,146 to 6,473 | 4,982 to 5,364 | 1,867 to 2,113 | 1,621 to 1,854 |
| Y-intercept when X=0.0 | 0,9679 to 1,187 | 0,8290 to 1,074 | 0,8016 to 0,9588 | 0,9651 to 1,115 |
| X-intercept when Y=0.0 | -0,1918 to -0,1505 | -0,2135 to -0,1560 | -0,5071 to -0,3841 | -0,6795 to -0,5266 |
| **Goodness of Fit** | | | | |
| R square | 0,976 | 0,9722 | 0,9262 | 0,9138 |
| Sy.x | 0,488 | 0,4182 | 0,2685 | 0,2552 |
| **Data** | | | | |
| Number of X values | 21 | 21 | 21 | 21 |
| Maximum number of Y replicate | 10 | 7 | 7 | 7 |
| Total number of values | 142 | 85 | 85 | 85 |
| Number of missing values | 68 | 125 | 125 | 125 |
| Equation | Y = 6,310*X + 1,077 | Y = 5,173*X + 0,9515 | Y = 1,990*X + 0,8802 | Y = 1,738*X + 1,040 |

| Linear regression | $N^{CTD}$+ATP | $N^{CTD}$+CTP | $N^{CTD}$+UTP |
|---|---|---|---|
| **Best-fit values** | | | |
| Slope | 6,593 ± 0,1503 | 5,725 ± 0,1087 | 6,163 ± 0,1606 |
| Y-intercept when X=0.0 | 0,9751 ± 0,1064 | 1,127 ± 0,07695 | 0,8585 ± 0,1101 |
| X-intercept when Y=0.0 | -0,1479 | -0,1969 | -0,1393 |
| 1/slope | 0,1517 | 0,1747 | 0,1623 |
| **95% Confidence Intervals** | | | |
| Slope | 6,292 to 6,894 | 5,507 to 5,943 | 5,840 to 6,485 |
| Y-intercept when X=0.0 | 0,7618 to 1,188 | 0,9728 to 1,281 | 0,6377 to 1,079 |
| X-intercept when Y=0.0 | -0,1866 to -0,1118 | -0,2305 to -0,1653 | -0,1822 to -0,09972 |
| **Goodness of Fit** | | | |
| R square | 0,9722 | 0,9805 | 0,9646 |
| Sy.x | 0,5817 | 0,4209 | 0,5997 |
| **Data** | | | |
| Number of X values | 19 | 19 | 19 |
| Maximum number of Y replicate | 3 | 3 | 3 |
| Total number of values | 57 | 57 | 56 |
| Number of missing values | 153 | 153 | 154 |
| Equation | Y = 6,593*X + 0,9751 | Y = 5,725*X + 1,127 | Y = 6,163*X + 0,8585 |

**Supplementary Table III: Comparison of quenching slopes.**

| ANOVA table | SS | DF | MS | F (DFn, DFd) | P value |
|---|---|---|---|---|---|
| Treatment (between columns) | 159,4 | 6 | 26,56 | F (6, 33) = 527,0 | P < 0,0001 |
| Residual (within columns) | 1,663 | 33 | 0,0504 | | |
| Total | 161 | 39 | | | |

| Tukey's multiple comparisons test | Mean Diff, | 95% CI of diff, | Significant | Summary | Adjusted P Value |
|---|---|---|---|---|---|
| $N^{CTD}$ vs. $N^{CTD}$+GTP | 1,137 | 0,7899 to 1,484 | Yes | **** | < 0,0001 |
| $N^{CTD}$ vs. $N^{CTD}$+ATP | -0,283 | -0,7466 to 0,1806 | No | ns | 0,4852 |
| $N^{CTD}$ vs. $N^{CTD}$+CTP | 0,585 | 0,1214 to 1,049 | Yes | ** | 0,0063 |
| $N^{CTD}$ vs. $N^{CTD}$+UTP | 0,147 | -0,3166 to 0,6106 | No | ns | 0,9516 |
| $N^{CTD-W330A}$ vs. $N^{CTD-W330A}$+GTP | 0,252 | -0,1245 to 0,6285 | No | ns | 0,3761 |

| Test details | Mean 1 | Mean 2 | Mean Diff, | SE of diff, | n1 | n2 | q | DF |
|---|---|---|---|---|---|---|---|---|
| $N^{CTD}$ vs. $N^{CTD}$+GTP | 6,31 | 5,173 | 1,137 | 0,1106 | 10 | 7 | 14,53 | 33 |
| $N^{CTD}$ vs. $N^{CTD}$+ATP | 6,31 | 6,593 | -0,283 | 0,1478 | 10 | 3 | 2,708 | 33 |
| $N^{CTD}$ vs. $N^{CTD}$+CTP | 6,31 | 5,725 | 0,585 | 0,1478 | 10 | 3 | 5,598 | 33 |
| $N^{CTD}$ vs. $N^{CTD}$+UTP | 6,31 | 6,163 | 0,147 | 0,1478 | 10 | 3 | 1,407 | 33 |
| $N^{CTD-W330A}$ vs. $N^{CTD-W330A}$+GTP | 1,99 | 1,738 | 0,252 | 0,12 | 7 | 7 | 2,97 | 33 |

**Supplementary Table IV: Statistical data of DSF and MST experiments.**

| **Boltzmann Sigmoidal fitting** | **Tm ± S.E.** | **Slope ± S.E.** | **$R^2$** |
|---|---|---|---|
| $N^{CTD}$ | 48.96 ± 0.03 | 0.79 ± 0.03 | 0.99 |
| $N^{CTD}$ + GTP | 49.97 ± 0.03 | 0.81 ± 0.02 | 0.99 |
| $N^{CTD-W330A}$ | 47.98 ± 0.07 | 2.18 ± 0.07 | 0.99 |
| $N^{CTD-W330A}$ + GTP | 47.67 ± 0.07 | 2.07 ± 0.07 | 0.99 |

| **MicroScale Thermophoresis** | **$N^{CTD}$** | **$N^{CTD-W330A}$** |
|---|---|---|
| Target Concentration: | 20nM | 20nM |
| Ligand Name | GTP | GTP |
| Ligand Concentration | 50 mM to 0.00153 mM | 50 mM to 0.00153 mM |
| n: | 3 | 3 |
| Excitation Power: | 77% | 77% |
| MST Power: | 40% | 40% |
| Temperature: | 25ºC | 25ºC |
| Kd (M): | 0.00019642 | 0.00085815 |
| Kd Confidence: | [0.00012558 - 0.00030723] | [0.00057568 - 0.0012792] |
| Response Amplitude: | 13.131 | 13.135 |
| Unbound: | 939.97 | 904.66 |
| Bound: | 926.84 | 899.98 |
| St. Error of Regression | 0.981 | 0.853 |
| Signal to noise: | 14.367 | 16.538 |
